# Lfc/Arhgef2 regulates mitotic spindle orientation in hematopoietic stem and progenitor cells and is essential for productive hematopoiesis

**DOI:** 10.1101/855577

**Authors:** Derek C. H. Chan, Ana Vujovic, Joshua Xu, Victor Gordon, Nicholas Wong, Laura P. M. H. de Rooij, Cailin E. Joyce, Jose La Rose, María-José Sandí, Bradley W. Doble, Carl D. Novina, Robert K. Rottapel, Kristin J. Hope

**Affiliations:** Stem Cell and Cancer Research Institute, Department of Biochemistry and Biomedical Sciences, McMaster University, Hamilton, Ontario, Canada; Michael G. DeGroote School of Medicine, Faculty of Health Sciences, McMaster University, Hamilton, Ontario, Canada; Department of Cancer Immunology and Virology, Dana-Farber Cancer Institute, Boston, Massachusetts, USA; Department of Medicine, Harvard Medical School, Boston, Massachusetts, USA; Broad Institute of Harvard and MIT, Cambridge, Massachusetts, USA; Princess Margaret Cancer Centre, University Health Network, Toronto, Ontario, Canada; Department of Medical Biophysics, University of Toronto, Toronto, Ontario, Canada; St. Michael’s Hospital, Division of Rheumatology, Department of Medicine, University of Toronto, Toronto, Ontario, Canada; Department of Immunology, University of Toronto, Toronto, Ontario, Canada

## Abstract

How hematopoietic stem cells (HSCs) coordinate their divisional axis relative to supportive niche cells and whether or not their divisional orientation is important for stem cell-driven hematopoiesis is poorly understood. Single cell RNA sequencing data from patients with the inherited bone marrow failure Shwachman-Diamond syndrome (SDS) show that *ARHGEF2*, a RhoA-specific guanine nucleotide exchange factor (GEF) and determinant of mitotic spindle orientation, is one of a restricted group of genes specifically downregulated in SDS HSCs and multipotent progenitors. Here, we describe Lfc/Arhgef2 as an important regulator of hematopoiesis *in vivo*. Transplanted Lfc/Arhgef2^-/-^ bone marrow shows impaired hematopoietic recovery and a production deficit of long-term HSCs. These phenotypes cannot be explained by differences in numbers of transplanted HSCs, their cell cycle status, level of apoptosis, progenitor output or homing ability. Using live imaging of dividing hematopoietic stem and progenitor cells (HSPCs), we show an increased frequency of misoriented divisions in the absence of Lfc/Arhgef2. Functional ARHGEF2 knockdown in human HSCs also impairs their ability to regenerate hematopoiesis, culminating in significantly smaller hematopoietic xenografts. Together, these data provide evidence demonstrating a conserved role for Lfc/Arhgef2 in orienting HSPC division and suggest that HSCs divide in certain orientations to establish hematopoiesis, the loss of which leads to their exhaustion in a mechanism that may underlie certain bone marrow failure syndromes.

## Introduction

Stem and progenitor cells across diverse species and organ systems are known to use both symmetric and asymmetric modes of division to achieve balanced expansion and differentiation during development. During embryogenesis, hematopoietic stem cells (HSCs) emerge in response to an asymmetric signal within the hemogenic endothelium^1^ and transit through both the fetal liver and spleen before reaching and colonizing the perinatal bone marrow.^2^ HSCs rapidly proliferate in this niche before becoming mostly quiescent,^3^ allowing their more proliferative downstream progeny, which follow complex lineage pathways,^4–6^ to productively drive native hematopoiesis.^7–12^ Numerous regulators of HSC activity that are either cell-autonomous^13^ or produced extrinsically by the niche^2^ have been identified to date; however, little is known about how HSCs divide and how their cell fate decisions are coupled to a divisional axis when establishing hematopoiesis in either native or transplant settings.

Dividing stem cells in other tissue types are known to couple their cell polarity axis with a properly oriented mitotic spindle to allow for appropriate acquisition of intracellular stem versus commitment fate determinants among daughter cells.^14^ Orientation of the mitotic spindle relative to stem cell-supportive niche signals is therefore critical for establishing cell polarity as well as directing fate decisions through the placement of prospective daughter cells in niche-proximal versus niche-distal conformations. For example, neural precursors in the developing brain undergo an early expansion phase, during which fate decisions are largely symmetric as a result of divisions occurring parallel to the apical neuroepithelium in the ventricular zone.^15,16^ A switch occurs in the subsequent neurogenic phase when the divisions of these expanded precursors become oblique and/or perpendicular, leading to the production of a differentiated progenitor cell that further contributes to proper cortical neurogenesis and layering.^17^ Within the intestine, dividing crypt base columnar stem cells perpendicularly align their mitotic spindles to the apical lumen, generating asymmetric daughter cell fates, whereas dividing cells in positions higher up the crypt assume parallel orientations.^18^ Similarly, in early epidermal development perpendicular mitotic divisions are utilized in order to properly specify epithelial stratification and differentiation.^19^

Whether or not similar principles hold true for the hematopoietic system remains to be determined. In the zebrafish, hematopoietic stem and progenitor cells (HSPCs) are found anchored to a mesenchymal stromal cell and achieve asymmetry of cell fates when they divide in orientations that displace one daughter cell away from this niche.^20^ Mammalian adult hematopoietic precursors are also known to divide in specific ways depending on their surrounding signaling milieu; a pro-renewal environment promotes symmetric expansion and a pro-differentiation environment biases for asymmetric fates.^21–23^ We and others have shown that HSC maintenance and activity are influenced by several known intrinsic effectors of polarity establishment and asymmetrically associated cell fate determinants.^24–32^ However, in distinct cell types and species, not all such factors appear to have the same mechanism or degree of action as initially described in model organisms.^32,33^ Nonetheless, one study to date that supports the importance of proper spindle orientation regulation in mammalian HSCs involves Lis1/Pafah1b1, a dynein-binding microtubule capturing protein that was shown to orient HSPC divisions and influence the inheritance of cell fate determinants in both normal hematopoietic and leukemia contexts.^34^ In this study however, both the functional importance of LIS1/PAFAH1B1 in human HSPCs and its potential role in non-malignant hematopoietic disorders were not explored.

Here, we describe Lfc/Arhgef2, a RhoA-specific GEF and determinant of mitotic spindle orientation, as an important regulator of mammalian hematopoiesis *in vivo*. We present evidence that hematopoiesis driven in the Lfc/Arhgef2^-/-^ background heavily relies on LT-HSCs and primitive progenitors and that a productive deficit is present at these levels within the hematopoietic hierarchy. We validate this functional link in human HSC-derived xenografts and directly show that Lfc/Arhgef2 regulates spindle orientation in HSPCs. We further show that *ARHGEF2* is specifically downregulated in HSC and primitive progenitor subsets from bone marrow samples of patients diagnosed with Shwachman-Diamond syndrome (SDS). Our findings demonstrate the importance of mitotic spindle orientation in HSPC function and suggest that depletion of *ARHGEF2* in humans may contribute to clinical bone marrow failure.

## Materials and Methods

### Generating and characterizing Lfc/Arhgef2 knockout mice

Lfc/Arhgef2^fl/fl^ and Lfc/Arhgef2^-/-^ mice were generated as previously reported^49^. Briefly, a loxP site was introduced upstream and a loxP-flanked reverse oriented neomycin resistance cassette downstream of exon 2 at the Lfc/Arhgef2 locus. Properly targeted ESCs were injected into recipient blastocysts and transplanted into the uteruses of foster mothers. The resulting chimeric mice were bred to C57Bl/6 females to establish a colony. A subset was then bred with a CMV-Cre expressing mouse strain to remove the loxP flanked exon 2 to yield a disruptive frameshift that resulted in loss of both a functional gene transcript and protein. Both exon-disrupted mice and mice that did not undergo Cre-lox mediated recombination were backcrossed for at least 4 generations.

### Colony forming unit, proliferation, cell cycle and apoptosis assays

To test hematopoietic progenitor outputs, 1.2 x 10^4^ whole bone marrow (WBM) from Lfc/Arhgef2^fl/fl^ and Lfc/Arhgef2^-/-^ mice were plated in biological triplicate in MethoCult GF M3434 (STEMCELL Technologies). Colonies were enumerated/scored between 12-14 days in culture. Lin^-^Sca-1^+^c-Kit^+^ (LSK) HSPCs were sorted and cultured for up to 7 days in HyClone Dulbecco’s High Glucose Modified Eagle Medium (GE Healthcare Life Sciences) supplemented with 10% fetal bovine serum (FBS; Gibco), 100 ng/mL murine stem cell factor, 100 ng/mL murine thrombopoietin, 10 ng/mL murine interleukin-3 and 10 ng/mL murine interleukin-6. K562 cells were cultured in IMDM (Thermo Fisher) supplemented with 10% FBS (Wisent Bioproducts) and 100 U/mL Pen/Strep (Thermo Fisher). Transductions were carried out by adding lentiviruses at a multiplicity of infection of 5 in the presence of 5 μg/mL polybrene (Sigma) for 3 days prior to *in vitro* assays. For cell cycle analyses, cells were fixed with Cytofix/Cytoperm solution (BD), permeabilized with Perm/Wash buffer (BD Biosciences) and stained with Ki-67-PE-Cy7 (BD) and 7-AAD (BD) to measure cells in G1 and S/G2/M phases. Early and late apoptosis were measured through combinatorial staining of Annexin V-FITC (BD) in binding buffer (BioLegend) and 7-AAD.

### Mouse bone marrow and fetal liver transplantation experiments

Non-competitive hematopoietic transplants were carried out in lethally irradiated (1100 cGy, Gammacell 40 Exactor, Best Theratronics) 8-to 12-week-old B6.SJL recipient mice. 1 x 10^6^ WBM cells or 3 x 10^5^ E14.5 fetal liver (FL) cells from Lfc/Arhgef2^fl/fl^ or Lfc/Arhgef2^-/-^ CD45.2+ donor mice or embryos were injected via tail vein. Competitive transplants were carried out using the same parameters but involved injecting either a 1:1 or 2:1 mixture of Lfc/Arhgef2^-/-^:C57Bl/6 WBM cells. Donor engraftment levels were serially monitored by tail vein blood collection and flow cytometry analysis using antibodies described below. Bone marrow transplant periods were ≥16 weeks in duration; secondary transplants were performed with doses of 1.5 x 10^6^ primary WBM cells. Kaplan-Meier survival analysis was calculated on the noncompetitive transplant cohorts. Homing experiments were conducted by injecting 5 x 10^4^ Lin^-^ cells into lethally irradiated recipients and re-isolating recipient bone marrow 16 hours later for flow cytometry analysis.

### Isolation of primary human HSCs and flow cytometry

Patient samples were obtained with informed consent and approved by the research ethics board at McMaster University. Umbilical cord blood cells were collected, lineage depleted and flow cytometrically analyzed as previously published.^33^ Antibodies against mouse antigens included: CD45.2 v450 (BD); Lineage eFluor 450 (eBioscience), Alexa Fluor 700 (BioLegend); Sca-1 APC (eBioscience), PerCP-Cy5.5 (eBioscience); c-Kit PE-Cy7 (BD); CD150 PE (BioLegend), Pacific Blue (BD); CD48 FITC (BD), APC (eBioscience); and CD11b PE, APC (BD); Gr-1 APC-Cy7, FITC, PE (BD); B220 PE, APC (BD); TER-119 PE (BD); CD3 PE (BD); CD4 PE (BD); and CD8a APC-Cy7 (BD). Sorting was performed on a MoFlo XDP (Beckman Coulter), routine acquisition on a LSRII (BD) and flow analysis completed using FlowJo v10.0.7 and v10.4 (Tree Star Inc.).

### Lentiviral constructs, knockdown and overexpression validation, and virus production

Third generation shRNA sequences against human ARHGEF2 and SBDS were selected based on high sensor assay rankings^35^ and cloned into lentiviral vectors pZIP-SFFV-ZsGreen-Puro (TransOMIC Technologies) or pZIP-SFFV-tNGFR-Puro that was re-adapted to contain an optimized miR-E delivery scaffold.^35^ Validation of knockdown efficiency was performed on leukemia cell lines and measured at the RNA level using quantitative polymerase chain reaction (qPCR) with the following primers: *ARHGEF2* (left 5’-TACCTGCGGCGAATTAAGAT-3’, right 5’-AAACAGCCCGACCTTCTCTC-3 ‘; Roche Universal ProbeLibrary #22); SBDS (left 5’-TGGCCAACAGTTAGAAATCGT-3’, right 5’-TTCCAAAGAACCTTTGCCTTTA-3’; Roche Universal ProbeLibrary #38); *EIF4H* (left 5’-CGTGGATCCAACATGGATTT-3’, right 5’-GGAGTCGTGGTCTCTGTGCT-3’ Roche Universal ProbeLibrary #35); and ACTB (Assay ID: Hs01060665_g1, Thermo Fisher). Human ARHGEF2 cDNA (NM_001162384.1) was subcloned into the pUMG-LV5 lentiviral expression vector (a gift from Maria Mesuraca). Protein level validation was performed by western blot using antibodies against human ARHGEF2 (Abcam, ab201687), SBDS (Abcam, ab128946) and ACTB (Santa Cruz Biotechnology, sc-81178). Lentiviruses were produced by co-transfection of the corresponding lentiviral vector with viral packaging plasmids pMD2.G and psPAX2 (a gift from Didier Trono, Addgene #12259 and #12260) into the 293FT cell line (Fisher Scientific) and concentrated by ultracentrifugation.

### Western blotting

Whole-cell lysates were prepared by lysing cells in RIPA buffer (50 mmol/L NaCl, 1% NP-40, 0.5% DOC, 0.1% SDS, 50 mmol/L Tris pH 8.0, 1 mmol/L EDTA) with complete EDTA-free Protease Inhibitor Cocktail (Milipore Sigma) and quantified using the Bradford assay (Bio-Rad). Protein samples were normalized to NuPAGE LDS sample buffer (Thermo Fisher Scientific) with β-mercaptoethanol (Sigma-Aldrich) and boiled for 5 minutes at 95°C prior to electrophoresis. Protein was transferred onto Immobilon-FL PVDF membrane (EMD Millipore), blocked with 5% skim milk powder in TBS-T for 30 minutes at room temperature and then incubated overnight at 4°C with primary antibodies. Following membrane washing, secondary antibody IRDye 680RD goat-anti-rabbit (LI-COR Biosciences) was added for 1 hour at room temperature and then imaged with the Odyssey Classic Imager (LI-COR Biosciences).

### Cord blood infection and xenotransplantation experiments

Xenotransplant experiments involved lentiviral transductions of shRNAs against a luciferase control or ARHGEF2 into Lin^-^CD34^+^ sorted human cord blood HSPCs and subsequent intrafemoral injections into sublethally irradiated (315 cGy) NSG mice. 5 x 10^4^ CD34^+^ HSPCs were cultured in StemSpan SFEM II (STEMCELL Technologies) supplemented with 100 ng/mL human stem cell factor, 100 ng/mL human fms related tyrosine kinase 3, 20 ng/mL human thrombopoietin and 20 ng/mL human interleukin-6 for 16 to 20 hours. Lentivirus was then added at a multiplicity of infection of 100. Transduced cultures were kept another 3 days before gene transfer values were measured by flow cytometry and 1/5 of day 0 equivalent cultures were transplanted into each recipient NSG mouse. Bone marrow aspirates were taken from the opposite femur between 8 to 10 weeks post-transplant to monitor interim engraftment; bilateral iliac crests, femurs and tibias were processed to evaluate the bone marrow ≥16 weeks post-transplant using flow cytometry.

### Live cell imaging of mitotic spindle orientation

To fluorescently label cells for mitotic spindle imaging, both H2B-EGFP and mCherry-α-tubulin were respectively cloned from pLKO-H2B-EGFP (a gift from Daniel Schramek) and mCh-alpha-tubulin (a gift from Gia Voeltz, Addgene #49149) vectors into a MSCV retroviral backbone. These constructs were cotransfected with pGP1 and pHCMV-G packaging plasmids into 293GPG packaging cells. Viral supernatant was serially passaged onto the GP+E-86 packaging cell line and double positive EGFP^+^mCherry^+^ producer cells sorted for co-culture with mouse LSK HSPCs over 3 days in equally mixed HyClone DMEM (GE Healthcare Life Sciences) and StemSpan SFEM (STEMCELL Technologies) media supplemented with 10% FBS (Gibco) and the aforementioned murine cytokines. CD45.2^+^EGFP^+^mCherry^+^ cells were then sorted and freshly plated onto retronectin coated chambered glass bottom μ-slides (Ibidi, 80447 and 80827). Live cell fluorescence images were acquired using a scanning laser confocal microscope with a 60x oil immersion objective (Nikon Eclipse Ti2). 15 μm stacks were obtained over 21 steps at 0.7 μm per step. Images were processed using ImageJ2^36^ on the Fiji platform^37^ where they were cropped to focus on dividing cells and rotated such that the plane of division was parallel to the X-axis of the image. Once aligned, the angle of division was visualized using an orthogonal view of the XZ plane. The angle of division was measured between the X-axis and the uppermost centrosome with the lowermost centrosome being used as the vertex of the angle.

## Results

### *ARHGEF2* is significantly downregulated in HSCs within patients with SDS

Defects at the stem and progenitor level underlie aspects of the severe paucity of hematopoiesis characteristic in inherited bone marrow failure syndromes.^38–44^ In exploring possible molecular mechanisms underpinning these disorders, we analyzed a recently published dataset of single cell RNA sequencing on CD34^+^ bone marrow HSPCs of patients diagnosed with SDS^45^ and observed that *ARHGEF2* was downregulated in SDS HSC/MPPs (FDR = 0.0029) and common myeloid progenitors (CMPs) (FDR = 0.0087) when compared to the same cell subsets found in healthy individuals (Figure 1A-C). *ARHGEF2* represented 1 of 229 genes found to be significantly and differentially reduced in expression in the HSC/MPPs of patients with SDS out of approximately 11,000 detected genes. Previous reports have demonstrated Lfc/Arhgef2 as a unique GEF in that it associates with, helps assemble and orients the mitotic spindle.^46–48^ In the mammalian neural system, downregulation of Lfc/Arhgef2 led to impaired neurogenesis and maintained precursors in a cycling state.^48^ In cord blood (CB) CD34+ cells, capitalizing on shRNA-mediated knockdown of SBDS, a known pathogenic driver in SDS, we show that the *ARHGEF2* transcript levels mirror the reduction in *SBDS* mRNA (Figure 1D, E). Moreover, while overexpression of ARHGEF2 did not alter early apoptosis levels in K562 cells targeted with shLuciferase, a trend of enhanced cell survival at the day 3 time point was observed with SBDS knockdown and ectopic ARHGEF2 expression, suggesting that ARHGEF2 may be dependent on and/or an important effector of disrupted SBDS activity (Figure 1D, F). Based on these combined data, we hypothesized that as it does in neural progenitors, Lfc/Arhgef2 similarly regulates HSC function and division within the hematopoietic system.

**Figure 1.**
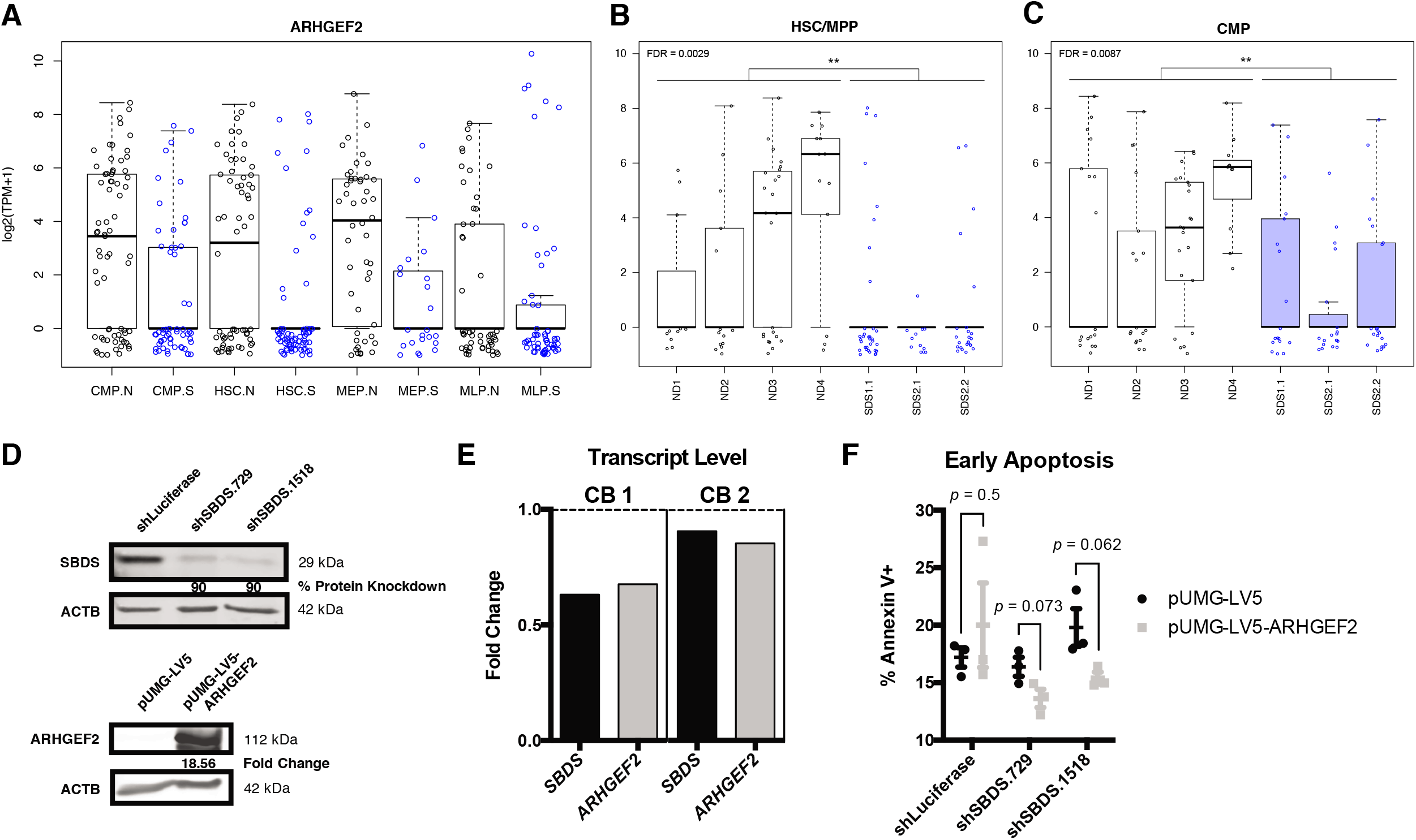
*ARHGEF2* is one of the most significantly downregulated genes in HSCs within patients with SDS. (A) *ARHGEF2* transcript expression within single hematopoietic stem cells (HSC), common myeloid progenitors (CMP), megakaryocyte-erythroid progenitors (MEP) and multi-lymphoid progenitor (MLP) cells from CD34^+^ patient samples from normal (.N, black) or Shwachman-Diamond syndrome (.S, blue) backgrounds. (B–C) *ARHGEF2* transcript expression from individual patient samples within HSC/MPP (B) (**, FDR=0.0029) and CMP (C) (**, FDR=0.0087) populations. ND and SDS represents normal donor and Shwachman-Diamond syndrome samples respectively. Each data point represents a single cell of each respective category. (D) *(Top)* Western blot validation of shRNAs targeting SBDS and *(Bottom)* ARHGEF2 overexpression. (E) Assessment of *SBDS* and *ARHGEF2* expression in human CD34^+^ HSPCs following SBDS knockdown as measured by qPCR (n = 2 cord blood samples). (F) Flow cytometric evaluation of early apoptosis levels upon concomitant ARHGEF2 overexpression and SBDS knockdown in K562 cells; n = 3 biological replicates evaluated at day 3 of *in vitro* cultures.

### Lfc/Arhgef2^-/-^ mice undergo compromised embryonic development and exhibit mild hematopoietic alterations at native steady-state

To directly investigate the effects of loss of function of Lfc/Arhgef2 in mammalian hematopoiesis, we capitalized on a previously validated Lfc/Arhgef2^-/-^ mouse model^49^ to first characterize the nature of steady state hematopoiesis in its absence (Figure 2A). While Lfc/Arhgef2^-/-^ mice were viable, litter sizes were noticeably reduced and serial heterozygous crosses yielded significantly fewer Lfc/Arhgef2^-/-^ mice than expected (Figure 2B, *Top*). In developing embryos, we observed an overall decrease in the percentage and absolute number of Lfc/Arhgef2^-/-^ fetal liver HSCs relative to controls (Figure 2B, *Bottom*). Among viable adult Lfc/Arhgef2^-/-^ mice, native peripheral blood analysis revealed approximately 25% fewer circulating platelets (Figure 2C), but no other significant differences in the number of leukocytes. These parameters were unchanged in both younger (4 months) and maturing adult (8 months) mice. Immunophenotyping of Lfc/Arhgef2^-/-^ adult bone marrow revealed a higher myeloid-to-lymphoid ratio that was maintained at the terminal end of the hierarchy (Figure 2D). There were significantly fewer lineage-negative cells (Figure 2E), characterized by fewer restricted (Lin^-^CD150^-^CD48+) and lymphoid (Lin^-^Sca-1^+^c-Kit^-^, LS) progenitors (Figure 2F) and a corresponding relative increase in myeloid (Lin^-^Sca-1^-^c-Kit^+^, LK) progenitors within the lineage-negative compartment (Figure 2F). However, neither HSPCs (Lin^-^Sca-1^+^c-Kit^+^, LSK) nor longterm HSCs (LSK CD150^+^CD48^-^, LT-HSCs) were significantly different between knockout and control adult bone marrow samples (Figure 2G). Overall, these data indicate that while Lfc/Arhgef2^-/-^ embryonic development and phenotypically primitive fetal HSC output is compromised and signs of thrombocytopenia are present in viable Lfc/Arhgef2^-/-^ mice, adult steady-state hematopoiesis is stable and only mildly altered in mice where the blood system is sufficiently established.

**Figure 2.**
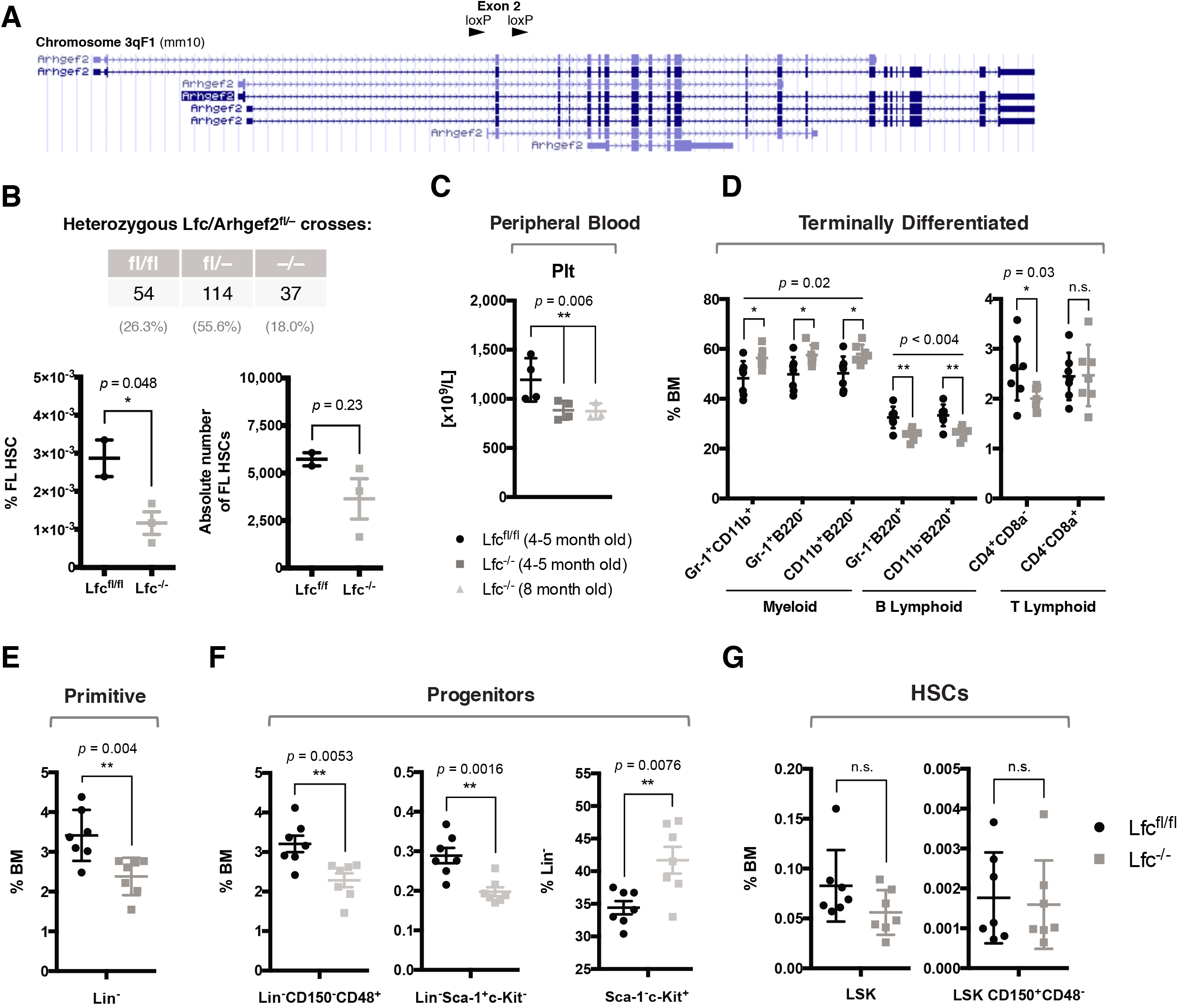
Native Lfc/Arhgef2^-/-^ mice exhibit mildly altered hematopoietic parameters. (A) Genomic locus of murine Lfc/Arhgef2 annotated with exon 2 flanked with loxP sites. (B) (*Top*) Non-Mendelian ratios observed from heterozygous Lfc/Arhgef2^fl/-^ crosses across 205 born pups. (*Bottom*) Relative percentage and absolute number of fetal liver HSCs (Lin^-^CD150^+^CD48^-^CD11b^+^) are decreased in Lfc/Arhgef2^-/-^ embryos. (C) Decreased circulating platelets in Lfc/Arhgef2^-/-^ mice (n = 4 mice per age group). (D) Higher myeloid-to-B lymphoid ratios, (E) fewer lineage-negative cell populations and (F) (*Left*) less restricted (*Middle*) lymphoid progenitors in Lfc/Arhgef2^-/-^ bone marrow. (F) (*Right*) Relative increase in myeloid progenitors within lineage-negative compartment of Lfc/Arhgef2^-/-^ bone marrow. (G) (*Left*) LSK HSPCs and (*Right*) LSK^+^SLAM LT-HSCs are not statistically different in Lfc/Arhgef2^-/-^ mice. (D–G) n = 7 mice per group. (C–G) * *p* < 0.05, ** p < 0.01; error bars represent standard error of the mean.

### Lfc/Arhgef2^-/-^ bone marrow HSPCs do not show significant alterations in their total colony output, proliferation, cell cycle or apoptosis status

To measure progenitor outputs, we performed colony forming unit (CFU) assays of whole bone marrow. While we did notice a slight decrease in the proportion of CFU-G, the remainder of all myeloid progenitors including CFU-GEMMs were present in similar numbers in Lfc/Arhgef2^-/-^ bone marrow as compared to Lfc/Arhgef2^fl/fl^ controls (Figure 3A). When measured at two distinct time points in culture, Lfc/Arhgef2^-/-^ LSK HSPCs did not differ in their proliferation rates (Figure 3B, *Left*), cell cycle status (Figure 3B, *Middle, Right*), or levels of early and late apoptosis (Figure 3C). These results argue against severe defects in either spindle stability (e.g. lack of and/or multipolar spindles) and DNA damage (e.g. aneuploidy, chromatin bridges) in these cells and show that apart from defects in granulocyte colony numbers, Lfc/Arhgef2^-/-^ HSPCs are functionally comparable to control HSPCs in their overall myeloid progenitor outputs, division kinetics, apoptosis and growth.

**Figure 3.**
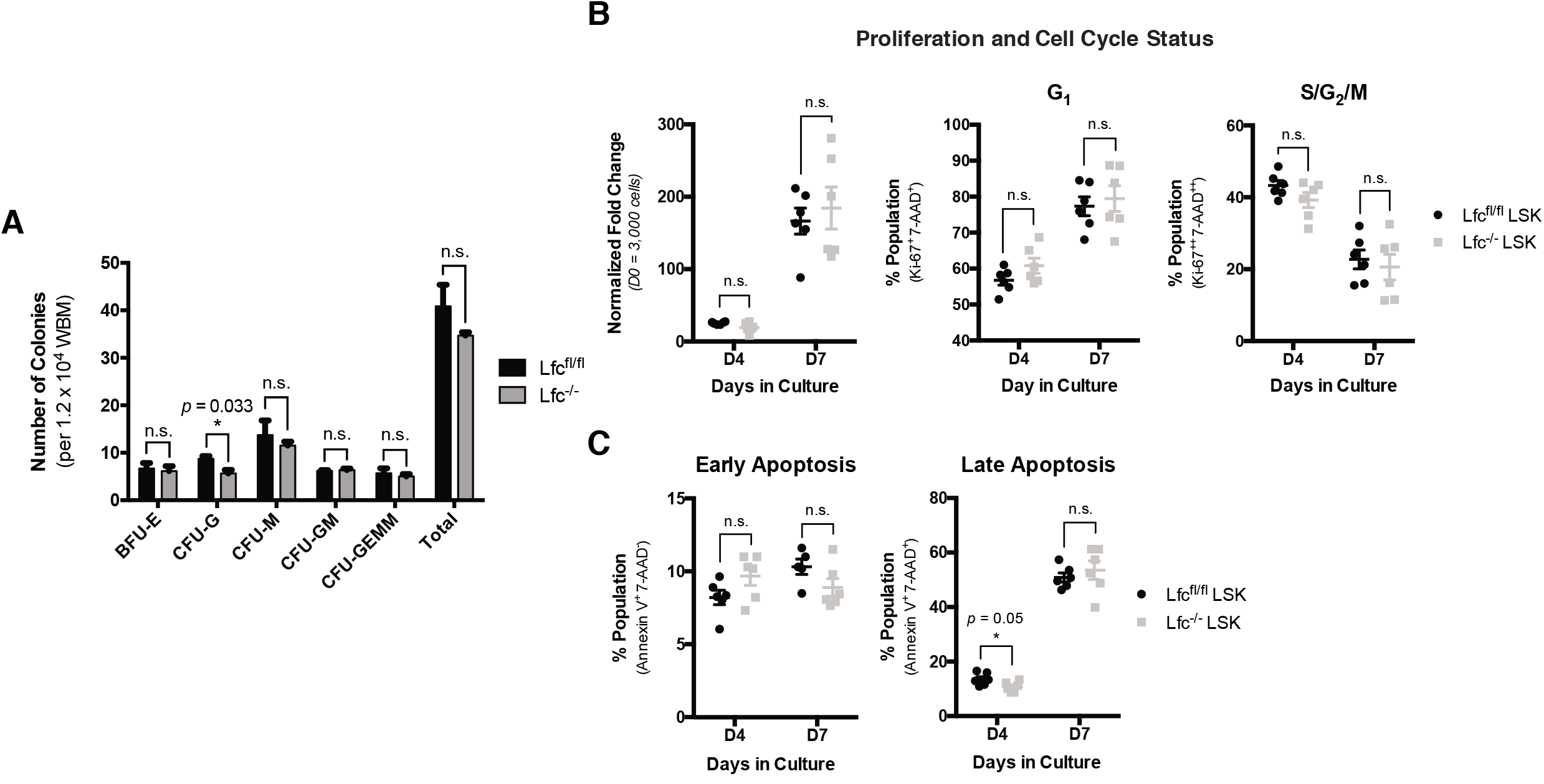
Lfc/Arhgef2^-/-^ bone marrow HSPCs do not show significant alterations in their total colony output, proliferation, cell cycle or apoptosis status. (A) Myeloid colony-forming units from 1.2 x 10^4^ whole bone marrow cells plated in biological triplicate enumerated at 7–10 days; error bars represent standard error of the mean; (B) (*Left*) LSK HSPCs cultured *in vitro* enumerated for proliferation, (*Middle*) proportional G1 and (*Right*) S/G2/M cell cycle status; and (C) (*Left*) early and (*Right*) late apoptosis at days 4 and 7 in culture. (B–C) n = 6 biological replicates; error bars represent standard error of the mean.

### Lfc/Arhgef2^-/-^ fetal liver and bone marrow insufficiently reconstitute the blood system, more heavily relies on and shows production deficits in HSCs

We next sought to verify that the decrease of phenotypic fetal liver HSCs was also apparent at the functional level by transplanting matched doses of E14.5 fetal liver cells isolated from Lfc/Arhgef2^fl/fl^ and Lfc/Arhgef2^-/-^ embryos into lethally-irradiated congenic recipients (Figure 4A). Within two weeks, the vast majority of recipients of Lfc/Arhgef2^-/-^ cells became moribund, while all mice having received Lfc/Arhgef2^fl/fl^ cells survived until the experimental 16-week post-transplant endpoint (Figure 4B). In 2 of the 6 Lfc/Arhgef2^-/-^ recipients that survived until 10 days post-transplant, we noted a relative decrease in the percentage of peripheral CD45.2^+^ and Gr-1^+^ granulocytic engraftment, with the lowest engrafted of these becoming moribund shortly after this sampling (Figure 4C). Together, this data indicates a significant impairment in functional repopulating HSCs within the fetal liver of Lfc/Arhgef2^-/-^ mice.

**Figure 4.**
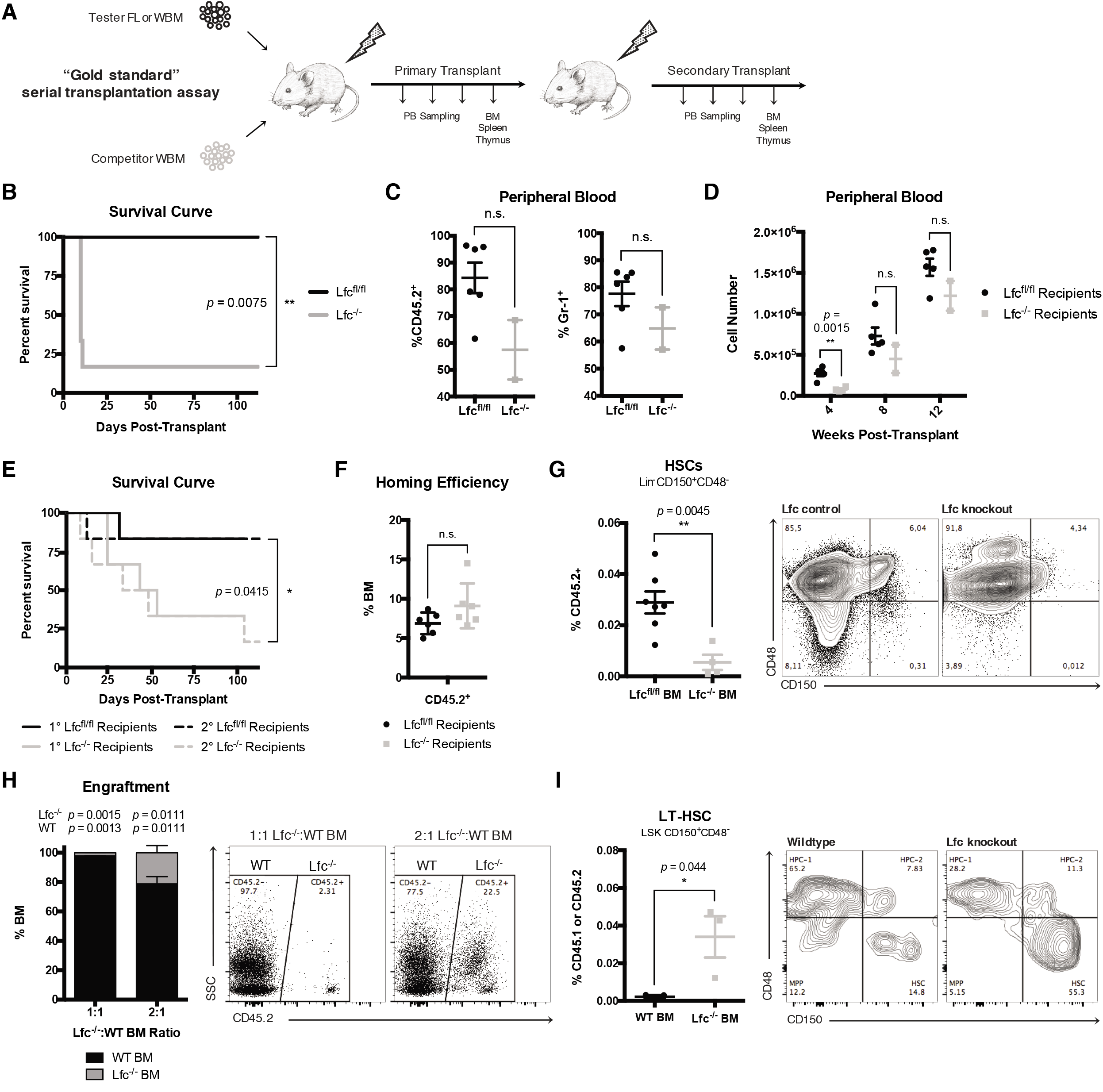
Lfc/Arhgef2^-/-^ fetal liver and bone marrow insufficiently reconstitute the blood system and show productive deficits at the HSC level. (A) Experimental schematic of non-competitive (test fetal liver or bone marrow only) and competitive (test and wildtype bone marrow) transplantations each lasting 10–16 weeks in duration. (B) Kaplan-Meier survival curves demonstrating significantly higher mortality among recipients of Lfc/Arhgef2^-/-^ E14.5 fetal liver cells; n = 6 recipients per experimental arm with n = 2 biological Lfc/Arhgef2^fl/fl^ and n = 3 Lfc/Arhgef2^-/-^ fetal liver donors. (C) Decreased levels of engraftment and granulocytic populations as measured in the peripheral blood of two remaining recipient mice of Lfc/Arhgef2^-/-^ E14.5 fetal liver cells at Day 10 post-transplant. (D) Insufficient and/or delayed hematopoietic recovery in the peripheral blood of primary transplanted mouse recipients of Lfc/Arhgef2^-/-^ bone marrow. (E) Kaplan-Meier survival curves demonstrating significantly higher mortality among recipients of noncompetitively transplanted Lfc/Arhgef2^-/-^ bone marrow. (F) Comparable homing efficiencies from 5 x 10^4^ Lin^-^ Lfc/Arhgef2^-/-^ bone marrow cells 16 hours after transplantation. (G) Significantly lower to near-absent levels of Lfc/Arhgef2^-/-^ Lin^-^CD150^+^CD48^-^ HSCs in secondary non-competitively transplanted bone marrow grafts. (H) Significantly poorer engraftment levels than expected from competing both 1:1 and 2:1 Lfc/Arhgef2^-/-^:wildtype doses of bone marrow among primary recipients. (I) Significantly increased proportion of LSK CD150^+^CD48^-^ LT-HSCs within secondary grafts among competitively transplanted recipients. (D–E) Primary non-competitive transplant arms were initiated each with n = 6 recipients using donor bone marrow derived from n = 3 biological replicates from either Lfc/Arhgef2^fl/fl^ or Lfc/Arhgef2^-/-^ mice. (F) Homing experimental arms each involved n = 6 recipients using bone marrow derived from n = 3 biological donor mice. (G) Secondary non-competitive transplant arm comparisons were completed with n = 7 Lfc/Arhgef2^fl/fl^ and n = 4 Lfc/Arhgef2^-/-^ recipient mice analyzed from n = 2 independent experiments. (H) Primary competitive transplants were set up with n = 3 recipients for each competing dose and (I) secondary competitive analyses were conducted with n = 3 recipients carried over from a total of n = 2 primary recipient donor mice. (B–I) * *p* < 0.05, ** p < 0.01; error bars represent standard error of the mean.

To functionally test the hematopoietic reconstitution capacity of Lfc/Arhgef2^-/-^ bone marrow, we performed competitive and non-competitive serial transplantation assays *in vivo* (Figure 4A). In non-competitive transplants, the majority of mice receiving Lfc/Arhgef2^-/-^ bone marrow showed evidence of anemia and became moribund, whereas recipients of Lfc/Arhgef2^fl/fl^ bone marrow did not display signs of hematopoietic insufficiency and/or delayed recovery (Figures 4D, E). Similar phenotypes and increased mortality were also evident in secondary transplant settings (Figure 4E). Importantly, this post-transplant failure phenotype was not due to compromised homing abilities, since as early as 16 hours posttransplantation, Lfc/Arhgef2^-/-^ lineage-negative cells homed to recipient bone marrow with an efficiency comparable to Lfc/Arhgef2^fl/fl^ lineage-negative cells (Figure 4F). However, within the grafts of the recipients of Lfc/Arhgef2^-/-^ bone marrow that remained at the end of secondary transplants, Lin^-^ CD150^+^CD48^-^ HSCs were significantly exhausted in comparison to those found in control grafts (Figure 4G). Competitive primary transplants of Lfc/Arhgef2^-/-^ bone marrow against wildtype bone marrow further demonstrated significantly impaired reconstitution at both equivalent (1:1) doses and when biased (2:1) to give an advantage to Lfc/Arhgef2^-/-^ bone marrow (Figure 4H). Interestingly, LT-HSCs were found to be significantly overrepresented in the few remaining Lfc/Arhgef2^-/-^ cells compared to Lfc/Arhgef2^fl/fl^ controls within grafts of secondary transplants (Figure 4I), highlighting a clear production deficit of downstream cells at the most primitive level. These findings demonstrate that in the absence of Lfc/Arhgef2, transplant- driven hematopoiesis heavily relies on LT-HSCs and primitive progenitors and that functional deficits existing at these levels lead to reduced hematopoietic engraftment and a subsequent increased mortality within recipient mice.

### Lfc/Arhgef2^-/-^ HSPCs exhibit a significantly increased frequency of misoriented divisions

Since Lfc/Arhgef2 has been uniquely characterized to function by orienting the mitotic spindle, we used a previously published live cell imaging method to measure the angle of division of mouse HSPCs.^34^ LSK cells were labelled with H2B-EGFP and mCherry-α-tubulin and plated on retronectin covered chambered slides. By acquiring confocal z-stacks of dividing cells and generating orthogonal projections of cell division events at telophase, we verified that wildtype LSK HSPCs preferentially divide parallel (between 0 and 10°) to this underlying substrate (Figure 5A). However, while Lfc/Arhgef2^-/-^ LSK HSPCs also yielded parallel division events, we observed a significantly increased frequency of non-parallel angles that reached as high as 60° (Figure 5B). These results indicate that while altered divisional preferences do not largely compromise HSPC survival and division kinetics *in vitro*, the post-transplant failure phenotypes measured *in vivo* could potentially be explained by dysregulated fate decisions as a result of misoriented divisions. Our results are thus consistent with the concept that Lfc/Arhgef2 is essential for regulating HSC divisional orientation and effective lineage differentiation within their niche during the establishment of hematopoiesis.

**Figure 5.**
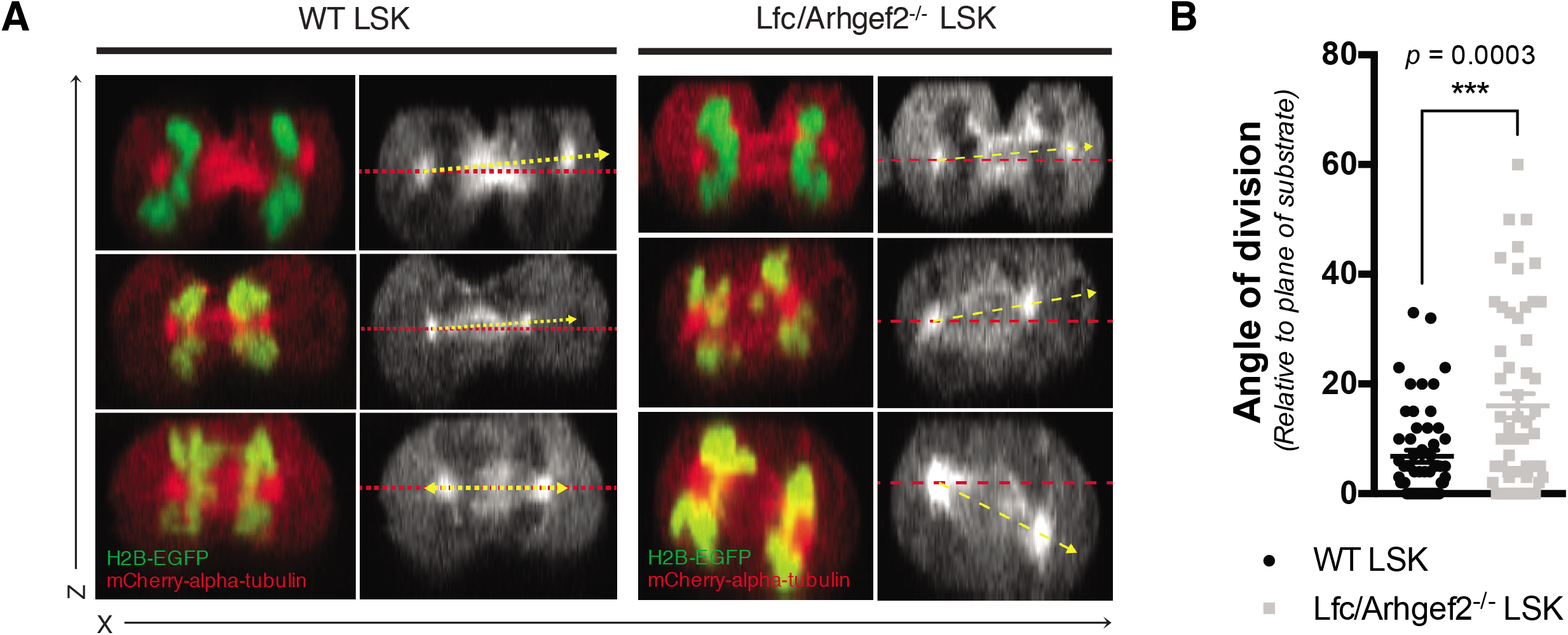
Loss of Lfc/Arhgef2 function disrupts HSPC mitotic spindle orientation. (A) Representative z-stack stitched images of LSK HSPCs retrovirally transduced with both H2B-EGFP and mCherry-α-tubulin imaged under live cell fluorescence microscopy to capture telophase events. (B) Quantification of cytokinesis events indicate that Lfc/Arhgef2^-/-^ LSK HSPCs exhibit a significantly increased frequency of random divisional orientations whereas wildtype HSPCs preferentially divide parallel to an underlying retronectin substrate. (n = 55 and 56 cells for wildtype and Lfc/Arhgef2^-/-^ backgrounds respectively).

### ARHGEF2 knockdown in human HSPCs compromises hematopoietic xenografts

To elucidate if ARHGEF2 function is conserved in human hematopoiesis, we performed immunofluorescence staining on several myeloid leukemia cell lines and confirmed that ARHGEF2 localizes at the microtubule apparatus during division (Figure 6A). To determine the functional effects of ARHGEF2 downregulation in human hematopoiesis, shRNAs against either ARHGEF2 or a luciferase control were introduced into cord blood CD34^+^ HSPCs (Figure 6B, C). Similar to results derived from Lfc/Arhgef2^-/-^ mice, cells with reduced ARHGEF2 proliferated comparably or was slightly dampened relative to controls (Figure 6D). Myeloid CFU assays yielded no significant differences in multipotent progenitor colonies and significantly decreased monocytic progenitors, while the total colony number remained equivocal across settings (Figure 6E). These data suggest that ARHGEF2 knockdown in human hematopoietic progenitors imparts only mild defects at the lineage-restricted level. Finally, using two separate and efficient shRNAs against ARHGEF2, *in vivo* analyses of intrafemorally xenotransplanted NSG recipient mice at 16 weeks post-transplant showed significantly diminished hematopoietic grafts with a paucity of CD15^+^ myeloid output observed in the residual xenografts of mice receiving ARHGEF2 knockdown cells compared to controls (Figure 6F, G). Considered with our murine data, this clear *in vivo* phenotype in the human context demonstrates the cross-species importance of ARHGEF2 to the regenerative and productive capacity of HSCs and may implicate ARHGEF2-regulated spindle orientation in human hematopoiesis (Figure 6H).

**Figure 6.**
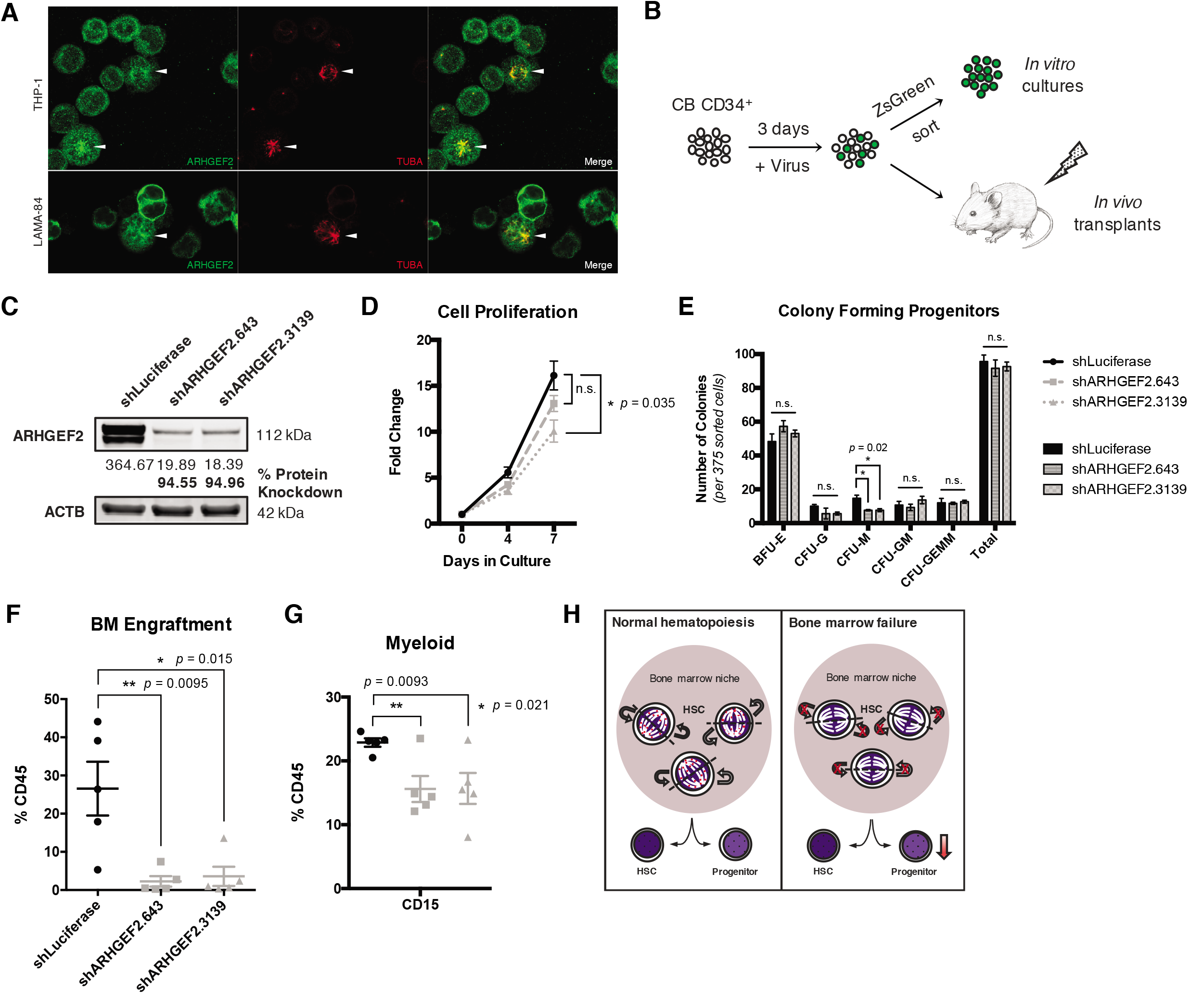
Loss of ARHGEF2 function in CD34^+^ HSCs results in significantly diminished xenografts. (A) Immunofluorescent staining showing co-localization of ARHGEF2 and TUBA at the mitotic spindle in human leukemia THP-1 and LAMA-84 cell lines. (B) Experimental schematic of shRNA knockdown of ARHGEF2 in CD34^+^ HSCs *in vivo* and *in vitro*. (C) Protein level knockdown validation of shRNAs against ARHGEF2. (D) Slightly decreased proliferation of CD34^+^ HSCs *in vitro* over 7 days. (E) Significantly fewer monocytic progenitor colonies but no differences in colony output overall in ARHGEF2 knockdown sorted cells. (F) Significantly decreased engraftments derived from CD34^+^ HSCs with comparable gene transfer levels receiving shRNAs targeting ARHGEF2 with significantly decreased outputs of CD15^+^ myeloid cells (G). Experimental arms included n = 5 recipients each derived from n = 1 cord blood sample. (H) Graphical model summarizing the role of Lfc/Arhgef2 (red dots) actively orienting the HSC mitotic spindle within the niche when establishing hematopoiesis, the loss of which leads to bone marrow failure at the stem cell level. (D–F) * *p* < 0.05, ** p < 0.01; error bars represent standard error of the mean.

## Discussion

In addition to its role in modulating RhoA activity at mitotic spindles,^46–48^ two studies to date have shown that Lfc/Arhgef2 associates with the microtubule array through the dynein light chain Dynlt1/Tctex-1^49–50^ and participates in a positive feedback loop in RAS transformed cells to potentiate MAPK signaling independent of its RhoGEF activity.^51^ While we cannot rule out the MAPK-regulatory function of Lfc/Arhgef2 underlying our observed phenotypes, the role of this pathway here is unlikely given that MAPK inhibition has been previously shown to have the opposite effect of improving HSC growth and output *in vitro* and *in vivo*.^52–54^ Global RhoA dependence has also been tested in the blood system however only in the context of conditional deletion within well-established chimeric grafts, where loss of the entire cellular pool of RhoA was found to not alter steady-state HSCs, but rather induce bone marrow failure due to significant progenitor loss.^55^ Our data using multiple transplant models highlights HSC but not pronounced progenitor deficiencies, suggesting that the regulation of RhoA activity at the mitotic spindle by Lfc/Arhgef2 represents an important axis for productive HSC divisions during the critical window over which hematopoiesis is established that may not be obvious at steady-state. In further support of this point, we note that examining the above-mentioned single cell RNA-sequencing data ARHGEF2, but not RHOA, is downregulated in primitive pediatric SDS cells (data not shown)^45^.

Our finding of decreased fetal liver HSC function in developing embryos of Lfc/Arhgef2^-/-^ mice indicates that defects in fetal hematopoiesis may also underlie the reduced fraction of Lfc/Arhgef2^-/-^ embryos that reach post-natal viability. Indeed, proper establishment of the hematopoietic system during development requires a minimum number of productive HSC divisions in the fetal liver.^56,57^ Thus, some but not all Lfc/Arhgef2^-/-^ embryos may generate enough effective HSC divisions to allow for sufficient downstream production of functional progenitors to populate the hematopoietic system. Our results *in vivo* using adult HSCs interrogated in two distinct transplant models provide further important insight into the mechanism of Lfc/Arhgef2 loss on HSC decision-making in different bone marrow states. In recipients of competitively transplanted Lfc/Arhgef2^-/-^ bone marrow cells, wildtype HSCs regenerated hematopoiesis more effectively, placing less of the reconstitution burden on Lfc/Arhgef2^-/-^ HSCs and allowing read-out of their preferred tendency to divide in a manner that promotes accumulation of primitive cells. This elevated HSC frequency may be due to an uncoupling of the cell polarity axis from their orientation of division, leading to a relative retention of stemness determinants in daughter cells. Alternatively, their compromised ability to adopt particular divisional orientations may result in Lfc/Arhgef2^-/-^ daughter cells being localized in more niche-proximal locations where they would receive enhanced HSC maintenance cues. In the non-competitive transplant setting, where in contrast the long-term regeneration of hematopoiesis is entirely dependent on Lfc/Arhgef2^-/-^ HSCs, the paucity of progenitors generated as a result of the spindle orientation defects leads to a heavier reliance on Lfc/Arhgef2^-/-^ HSC divisions. In this latter context, which importantly mimics the dependencies on HSCs in developmental hematopoiesis, the observed outcome of stem cell loss is likely due to an exhaustion of Lfc/Arhgef2^-/-^ HSCs.

Our observation of similarly defective hematopoietic reconstitution *in vivo* upon transplant of ARHGEF2- depleted human cord blood HSCs suggests a conserved function of Lfc/Arhgef2 across species. Downregulation of *ARHGEF2* in SDS HSCs and acutely upon SBDS repression in CD34^+^ cells points to the possibility that repression of ARHGEF2 in SDS patients may contribute to defective HSC-driven hematopoiesis. It is interesting to note that in our Lfc/Arhgef2^-/-^ mouse model we observe native thrombocytopenia, indicating an additional role for Lfc/Arhgef2 in regulating megakaryocyte maturation,^58,59^ defects in neutrophil chemotaxis,^60^ and while not formally characterized yet, bone malformations and clear neurological abnormalities (data not shown), the latter of which parallels reports of cognitive impairments and intellectual disability in patients with ARHGEF2 loss-of-function mutations.^61,62^ Importantly, all of these features can be found in patients diagnosed with SDS,^63,64^ which encourages future efforts to understand if the loss of ARHGEF2 function contributes to the etiology and pathogenesis of SDS.

Finally, our work highlights implications for how mitotic spindle orientation itself may more broadly affect stem cell division during development and disease. During brain development, centrosomal protein loss-of-function events that influence spindle orientation result in a disrupted balance of symmetric and asymmetric divisions that lead to microcephaly.^65^ Conceptual parallels may therefore also exist between spindle-regulating genes and bone marrow failure syndromes within and beyond SDS. Indeed, the loss of Cdk5rap2, a centrosomal spindle-orienting protein, results in a macrocytic, hypoproliferative anemia and leukopenia (“Hertwig’s anaemia”) in an irradiated mouse model.^66,67^ With the identification of several other spindle-regulating genes implicated in microcephaly, it may be interesting to determine if any of these genes also have roles within bone marrow failure or in disorders that result in tissue insufficiency elsewhere. Finally, spindle orientation dysregulation may also play an important role in diseases that potentially encourage enrichment of abnormal divisional axes within the niche and will therefore be interesting to explore as a possible contributor to disorders that include clonal hematopoiesis, myelodysplastic syndrome and/or leukemia.

## Acknowledgements

The authors thank Johann Hitzler and Sheila Singh for important feedback on this work; Minomi Subapanditha for flow cytometry sorting; Lillian Robson, Wendy Whittaker and Norma-Ann Kearns for animal caretaking and maintenance; and Ray Truant for access to the McMaster Biophotonics Facility.

This work was supported by a Canadian Institutes of Health Research (CIHR) MD/PhD Studentship (D.C.), Ontario Graduate Scholarship (D.C.), a CIHR PhD Studentship (J.X.), an Ontario Institute for Cancer Research (OICR) Investigator Award (K.H.) and a CIHR Foundation Grant (R.R.).

## Authorship

Contributions: D.C. designed, led and performed the experiments, analyzed and interpreted the data and wrote the manuscript. A.V., J.X. and L.d.R. assisted with animal experiments. V.G. and N.W. assisted with imaging work. C.J. and C.N. designed and completed analyses on sequencing data from patient samples. J.L.R., M.S., and R.R. generated the knockout mouse model. R.R. advised on experimental design. K.H. supervised the project, designed experiments, reviewed the data and wrote the manuscript. All authors reviewed and approved the manuscript.

